# Grape polyphenols reduce gut-localized reactive oxygen species associated with the development of metabolic syndrome in mice

**DOI:** 10.1101/331348

**Authors:** Peter Kuhn, Hetalben M. Kalariya, Alexander Poulev, David M. Ribnicky, Asha Jaja-Chimedza, Diana E. Roopchand, Ilya Raskin

## Abstract

High-fat diet (HFD)-induced leaky gut syndrome combined with low-grade inflammation increase reactive oxygen species (ROS) in the intestine and may contribute to dysbiosis and metabolic syndrome (MetS). Poorly bioavailable and only partially metabolizable dietary polyphenols, such as proanthocyanidins (PACs), may exert their beneficial effects on metabolic health by scavenging intestinal ROS. To test this hypothesis, we developed and validated a novel, noninvasive, in situ method for visualizing intestinal ROS using orally administered ROS-sensitive indocyanine green (ICG) dye. C57BL/6J mice fed HFD for 10 weeks accumulated high levels of intestinal ROS compared to mice fed low-fat diet (LFD). Oral administration of poorly bioavailable grape polyphenol extract (GPE) and β-carotene decreased HFD-induced ROS in the gut to levels comparable to LFD-fed mice, while administration of more bioavailable dietary antioxidants (α- lipoic acid, vitamin C, vitamin E) did not. Forty percent of administered GPE antioxidant activity was measured in feces collected over 24 h, confirming poor bioavailability and persistence in the gut. The bloom of beneficial anaerobic gut bacteria, such as *Akkermansia muciniphila*, associated with improved metabolic status in rodents and humans may be directly linked to protective antioxidant activity of some dietary components. These findings suggest a possible mechanistic explanation for the beneficial effects of poorly bioavailable polyphenols on metabolic health.

## Introduction

Incidence of metabolic syndrome (MetS) and type-2 diabetes (T2D) is rapidly growing in the world’s population [1], highlighting the importance of new approaches for prevention, detection, and treatment. Clinical and epidemiological studies as well as associated meta-analyses, suggest that consumption of diets rich in plant polyphenols offer protection against development of chronic non-communicable diseases (NCDs) such as MetS and T2D [2-5]. Studies have also shown that dietary polyphenols from a variety of fruits (blueberry, apple, cranberry, grape) can attenuate the symptoms of MetS in mice [6, 7] and most recently, these improvements have been associated with alterations in the gut microbiota [8-10]. However, the mechanism by which these fruit polyphenols, particularly large molecular weight oligomeric and polymeric flavonoids, exert their beneficial health effects remains uncharacterized. The hypothesis that dietary antioxidants, such as polyphenols, may protect against chronic NCDs was formulated more than 60 years ago [11]. This hypothesis gave birth to the widespread popularity of antioxidants found in fruits, vegetables, chocolate, coffee, tea, wine and contributed to the development of the dietary supplement industry.

Notably, antioxidant flavonoids, such as anthocyanins, monomeric flavan-3-ols, and their oligomers, B-type proanthocyanidins (PACs), which represent the majority of polyphenols in grapes [12-14] and other red- or blue-colored berries are poorly bioavailable [15-18]. Relatively low systemic absorption of these compounds, combined with their documented health benefits, creates a paradox in understanding the mechanism of action of berry polyphenols and leads to several hypotheses aimed at explaining their health benefits. Colonic microbes can metabolize berry flavonoids and some of the resulting metabolites have been suggested to be responsible for the observed health benefits [19, 20]. Studies from several laboratories have shown that grape, cranberry, and apple polyphenols promote the growth of mucin-degrading gut bacterium *Akkermansia muciniphila* in association with leaner phenotype, less intestinal and systemic inflammation, improved oral glucose tolerance, and intestinal gene expression consistent with improved gut barrier and metabolism [8-10]. Oral administration of *A. muciniphila* cultures in high-fat diet (HFD)-fed mice attenuated gut barrier dysfunction and other symptoms of MetS [21] while administration of Amuc_1100*, an outer membrane protein of *A. muciniphila* that interacts with TLR2, improved gut barrier and partially reproduced the beneficial effects of the bacterium in mice [22, 23]. Increased intestinal abundance of *A. muciniphila* has also been correlated with anti-diabetic and anti-obesity therapies such as metformin treatment [24, 25] and gastric bypass surgery [26], further supporting its positive impact on metabolic health.

While the ability of poorly bioavailable berry polyphenols to scavenge reactive oxygen species (ROS), has been widely associated with their health benefits, little is known about the presence and distribution of ROS or molecular oxygen in human gut. A steep oxygen gradient is assumed to exist between intestinal submucosa adjacent to the mesentery blood vessels to the center of the lumen, which is almost anoxic [27]. These assumptions are based on the relatively invasive measurements with polarographic Clark-type electrodes [28, 29] or on less invasive and sensitive electron paramagnetic resonance (EPR) oximetry [30]. Increased intestinal permeability, sometimes referred to as leaky gut syndrome, is usually associated with MetS, obesity, chronic low grade intestinal inflammation, and gut dysbiosis [31, 32]. It is manifested in a transport of water into the intestinal lumen and leakage of pro-inflammatory microbial lipopolysaccharide (LPS) into the bloodstream, which can promote low-grade, systemic inflammation and insulin resistance [33, 34]. Increased intestinal permeability may also be associated with greater diffusion of oxygen and ROS in the intestine from the mesenteric vasculature, which, in turn, may affect gut microbiome communities by favoring oxygen-tolerant bacteria (facultative anaerobes) at the expense of microbes that thrive under anaerobic or microaerophilic conditions, such as *A. muciniphila*. Inflammation, associated with inflammatory bowel disease (IBD) and obesity, also leads to the localized and systemic production of ROS [35, 36]. Therefore, pro-inflammatory disorders may lead to further increase in intestinal ROS and cause depletion of anaerobic bacterial species, which are particularly sensitive to ROS, as they lack biochemical defenses systems against their toxic effects [37].

Optical in vivo imaging using near-infrared fluorescence (NIRF) light generated by cyanine-based fluorescent dyes permits relatively deep photon penetration into tissue, minimal auto-fluorescence, less scatter, and high optical contrast [38, 39]. To better understand the impact of poorly bioavailable, antioxidant dietary polyphenols on ROS in vivo, we developed a non-invasive *in situ* method using cyanine-based dyes to measure intestinal abundance of ROS. In biological systems hydrocyanines selectively react with ROS, such as superoxide, via an amine oxidation mechanism to regenerate fluorescing cyanine dyes, thus allowing imaging of nanomolar levels of ROS. The resulting near infrared fluorescence (NIRF) is directly related to ROS content. Both cell-permeable and impermeable variants of hydrocyanines were developed and used to study surgically created ischemia [40], implant-associated inflammation [41], cancer development [42], and commensal bacteria-induced ROS production in intestinal epithelial sections [43, 44]. Cell-impermeable hydrocyanines were used successfully for the in vivo imaging of retinal oxidative stress in rats [45].

To the best of our knowledge, the present study is the first to use cell-impermeable hydrocyanines, specifically indocyanine green (ICG), to image ROS in the gut of live animals using non-invasive fluorescent imaging techniques. This manuscript reports on the development of ICG florescence-based ROS imaging methodology for the gut of live animals and the application of this methodology to study the effects of grape polyphenols and other dietary antioxidants on gut ROS content in healthy and HFD-fed, metabolically compromised animals. Our findings indicate that poorly absorbed dietary polyphenols effectively counteract obesity-associated ROS increase in the gut, and thus may initiate the cascade of events that lead to the improvement in MetS and associated dysbiosis.

## Materials and Methods

### Mice

Animal studies were conducted at an AAALAC-approved facility of Rutgers University using Rutgers IACUC-approved protocols. Twenty-five C57BL/6J diet-induce obese (DIO; 60 kcal% fat diet; Research Diet #D12492) and twenty-five C57BL/6J control (10 kcal% fat diet; Research Diets #D12450J) male mice (Jackson Labs, Cat# 380050) were purchased at 13 wk-old. Both sets of mice were randomly divided into five groups (1 control, 4 experimental groups) of five mice and maintained on their respective diets. Animals were acclimated for two weeks before experimentation and housed at a constant temperature on a 12 h light/dark cycle with free access to food and water.

### Phytochemicals and reagents

Indocyanine green (ICG), β-carotene, gallic acid, α-lipoic acid, L-ascorbic acid (vitamin C), D-alpha-tocopherol succinate (vitamin E) and 2,2’-azino-bis (3-ethylbenzothiazoline-6-sulphonic acid (ABTS), 6-hydroxy-2,5,7,8-tetramethylchroman-2-carboxylic acid (TROLOX), purified oligomeric B-type proanthocyanidins from grape seed (Cat# 1298219), and procyanidin B2 analytic standard (Cat # 29106-49-8) were purchased from Sigma-Aldrich, St Louis, MO. Grape polyphenol extract (GPE) was prepared from frozen grape pomace (provided by Welch’s, Concord, MA). Details of grape pomace extraction and column-purification, GPE biochemical characterization, and colorimetric quantification of total polyphenols and PAC contained in GPE are described in Supplementary Methods.

## Grape polyphenol extract (GPE)

Frozen grape pomace was added to 50% ethanol in a ratio of 1:5 (grams:milliliters) and thoroughly ground in a Vitamix^®^ Blender. The pH of the pomace slurry was decreased to 2.5 with concentrated sulfuric acid, and the mixture extracted for 2 h at 80°C with agitation. The extract was separated from the solids with a filtration centrifuge (Model RA-20VX, Rousselet Robatel Co., Annonay, France). The filtered extract was concentrated to 100 mL in a rotary evaporator and the polyphenol fraction further purified using solid-phase extraction (SPE) Strata C-18-E 20 g / 60 mL column (Phenomenex, Torrance, CA). The column was prewashed using 2 bed volumes each of ethyl acetate, followed by methanol acidified with 1% acetic acid, and then water acidified with 1% acetic acid. After loading the extract (approximately 20 ml) onto the column, the column was washed with 3 bed volumes of water acidified with 1% acetic acid. Thereafter, polyphenols were eluted with methanol acidified with 1% acetic acid. The eluate was collected and rotary-evaporated to dryness. To fully remove GPE from the evaporating flasks, extract was re-dissolved in water, freeze-dried, and stored −20°C until use.

## GPE characterization by LC-MS/MS and colorimetric quantification

Procyanidin B2 was used as an external standard for the quantification of some of the individual compounds, as procyanidin B2 equivalents exist in GPE and in PACs. We previously verified that PACs with different degrees of polymerization accounted for 90% of the purified grape seed PACs purchased from Sigma-Aldrich and used in our experiments and performed a detailed LC-MS analysis of the sample, using the methodology described in Zhang *et al.* [46]. PACs, quantified with DMAC assay adapted from Prior *et al*. 2010 [47] and total polyphenols, quantified with Folin-Ciocalteau assay [48] were found to comprise 56% and 69% of GPE, respectively.

Using this LC-MS method, we were able to quantify PAC monomers, dimers, trimers, tetramers and pentamers, as well as their corresponding gallates (**Supplementary Fig 1**). Overall, these compounds comprised 18% GPE. The discrepancy between colorimetric and LC-MS quantification was attributed to the presence of multiple PAC derivatives not quantified with the LC-MS method. The colorimetric method, on the other hand, quantifies total PACs and polyphenols in the sample. Qualitatively, the biochemical composition of GPE and the oligomeric B-type PAC sample from Sigma-Aldrich [46] was found to be similar.

### Feces extraction for antioxidant and total polyphenol assays

Fifteen wk-old mice were gavaged with GPE delivering 32 mg total polyphenol / kg of body weight. Feces were collected hourly from 1 h to 12 h and at 24 h after the gavage. Feces were mixed with 50% Ethanol (100 mg/1 mL), then homogenized with a Geno/Grinder (Model 2010, Metuchen, NJ) at 1500 rpm for 8 min or ultra-sonicated for 30 min. Samples were centrifuged at 13,000 rcf for 40 min. The supernatant was collected and used for 2,2’-azino-bis(3-ethylbenzothiazoline-6-sulphonic acid (ABTS) antioxidant assay [49] and Folin–Ciocalteu assay to quantify total polyphenols. For the ABTS assay, 7 mg/mL ABTS solution and 50 mg/mL K_2_S_2_O_8_solution were prepared in 1 mL of milliQ water. 20 µL 50 mg/mL K_2_S_2_O_8_were added to the ABTS solution before the mixture was incubated for 30 min and then diluted to 20 mL of MilliQ water. TROLOX standards were prepared in 95% ethanol from 600 µg/mL stock and diluted to a standard range. For the assay, 1 mL of ABTS was added to 50 µL of samples or standard, briefly vortexed, then loaded to a microwell plate in duplicate. Absorbance at 734 nm was read within 4 min using a BioTek Synergy HT Multi-Detection Plate Reader.

### Preparation of hydro-indocyanine green (H-ICG)

H-ICG used for the studies was prepared from the cyanine dye, indocyanine green, by reduction with NaBH_4_. Indocyanine green and its hydrocyanine product H-ICG are generally membrane impermeable, nontoxic and should remain in the gastrointestinal track until eliminated via feces. Briefly, 8 mg of dye was dissolved in 8 mL methanol and reduced by adding 4-8 mg of NaBH_4_. The reaction mixture was stirred for 5-10 minutes and solvent removed under reduced pressure. The resulting precipitate was nitrogen capped and stored at −20°C.

### In vivo ROS imaging and analyses

Prior to imaging, body hair around the abdomen and back were shaved and depilated using Veet^®^ hair remover followed by a water rinse. At 16 weeks of age, animals were gavaged with vehicle or GPE (total polyphenol dose of 32 mg/kg), PAC (32 mg/kg), β-carotene (32 mg/kg), or a mixture of L-ascorbic acid, D-α-tocopherol succinate, and α-lipoic acid (each 32 mg/kg). Mixing H-ICG with GPE reduced the sensitivity of the measurements, so H-ICG was reconstituted in water (1 mg/mL) and orally administered (dose of 6 mg/kg) 1 h after administration of the antioxidants, which resulted in the most reliable and reproducible fluorescent images (data not shown). In-Vivo MS FX PRO imaging system (Bruker, Ettlingen, German), equipped with Bruker Carestream Multimodal Animal Rotation System (MARS), was used to capture both brightfield and NIRF images of the experimental animals at precise and reproducible positioning angles. Mice were anesthetized with 2% isoflurane and placed in the imaging system 45 min after dye administration (1 h 45 min after polyphenol/antioxidant treatment). Isoflurane anesthesia (1-2%) was maintained during imaging procedure. A small bead of sterile artificial tears ointment was applied to each eye of mice to maintain lubrication.

Mice were initially placed into the MARS system in a supine position with their spine directed towards the camera. Each mouse was rotated at 30° increments over 360° for 24 min and fluorescence and brightfield images were taken consecutively at each angle. Rotation of each mouse started 45 min following dye administration so that imaging of ventral orientation corresponded with strongest signal, approximately 1 h after dye administration. Following excitation illumination at 760 nm, emission at 830 nm was recorded using a filter equipped high sensitivity cooled charged coupled device camera. Acquisition time was 20 s for each fluorescent image, followed by a brightfield light photograph (0.5 s exposure). Both NIRF and brightfield images were optically superimposed to visualize anatomical information.

Quantitative analysis of the optical signal capture was completed in Carestream MI software v5/0.529 (Carestream Health Inc.). Fluorescence intensity within a rectangular area of fixed dimensions (161 by 158 pixels) was recorded. Images from control mice (i.e. not gavaged) were used to subtract background and the mean fluorescence intensity for each image was determined. Fluorescence images were converted to photons/s/mm^2^ using Bruker imaging software. Data were presented with the fluorescence values as a function of the imaging angles.

### Statistical analysis

Statistical analyses were conducted using GraphPad Prism 5. Details are provided in figure legends.

## RESULTS

### Excretion of grape polyphenols and related antioxidant activity

Fecal samples from mice gavaged with GPE (32 mg total polyphenols/kg) showed a large increase in antioxidant activity (**Fig. 1**). This increase was apparent at 4 h in mice allowed *ad libitum* access to food after treatment (**Fig. 1A**) and at 3 h in mice fasted for 12 h after treatment (**Fig. 1B**). Fecal antioxidant activity in fed animals returned to basal levels 12 h after treatment, while elevated fecal antioxidant capacity could still be detected at 12 h in fasted mice, after which time *ad libitum* feeding was resumed. Fecal antioxidant activity of these initially fasted mice returned to basal levels by 24 h.

**Figure 1.**
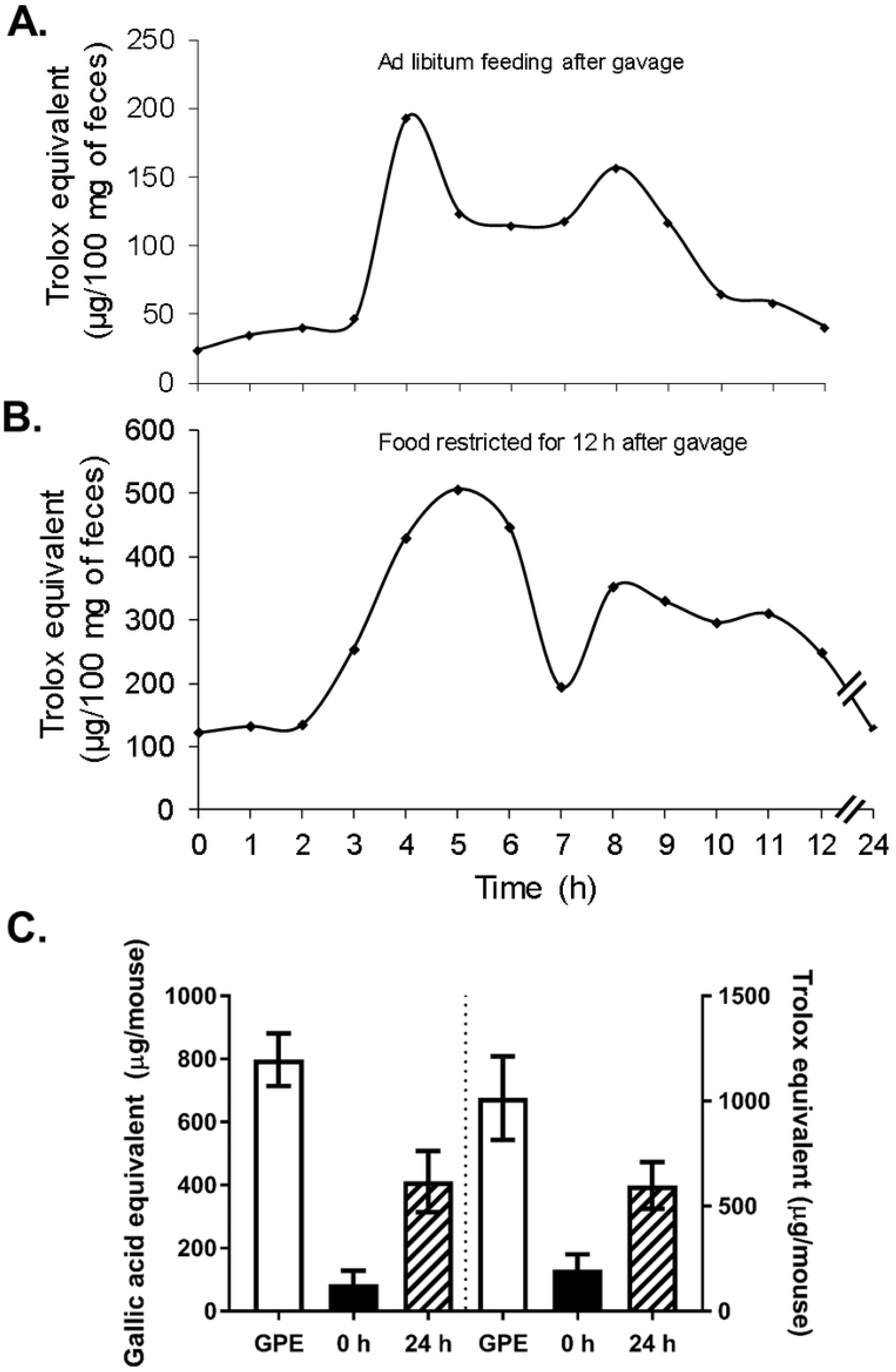
Fecal samples collected after GPE treatment contain high levels of antioxidant activity. Antioxidant activity (Trolox equivalents) in murine fecal samples (n= 6 mice) collected before (0 h) a single oral dose of GPE (32 mg total polyphenols/ kg) and **A.** every hour for 12 h after treatment while mice had *ad libitum* access to chow diet or **B.** every hour during a 12 h period of food restriction, after which chow diet was replaced and an additional fecal sample was collected at 24 h. **C.** Total polyphenols (as gallic acid equivalents, left) and total antioxidant capacity (as Trolox equivalents, right) in fecal samples collected before gavage (0 h, black bars) and in total feces collected over the 24 h period following oral GPE administration (crosshatched bars), as compared to the initial dose (white bars). N=6 mice; Data are reported as mean ± SD.

In a separate experiment, fecal samples were collected from mice before gavage (0 h) and total feces were collected over the 24 h period after gavage with GPE (66 mg GPE equivalent to 32 mg total polyphenols/ kg body weight). Fecal samples were pooled over a 24 h period and analyzed for antioxidant capacity (as Trolox equivalents) and total polyphenol content (as gallic acid equivalents; **Fig. 1C**). Based on the reactivity in Folin–Ciocalteu and ABTS assays, these data suggest that approximately 40% of the initially gavaged GPE antioxidant activity could be recovered in the feces.

### Imaging of reactive oxygen species (ROS) in mouse gut: timing and intensity

Mice fed both diets for 15 weeks were gavaged with GPE followed by H-ICG, anesthetized and imaged for NIRF at 830 nm (excitation 760 nm) in a ventral orientation (abdomen facing camera) 1, 2 and 3 h following H-IGG administration (2, 3 and 4 h following GPE administration). These experiments were designed to determine the timing of maximum ROS-induced fluorescence response following the administration of GPE and H-ICG and to compare levels of ROS in the guts of LFD-fed and HFD-fed mice. A significant and reproducible increase in ROS content was observed in HFD-fed mice compared to LFD-fed mice (**Fig. 2A**) at 1 h and 2 h measurements (**Fig. 2B**). The greatest ROS signal was observed 1 h after H-ICG administration (2 h after GPE administration) and decreased by 3 h following H-IGG administration (**Fig. 2B**). Based on these data, all subsequent in vivo ROS measurements were carried out as close as possible to 1 h following H-ICG administration (2 h following GPE administration).

**Figure 2.**
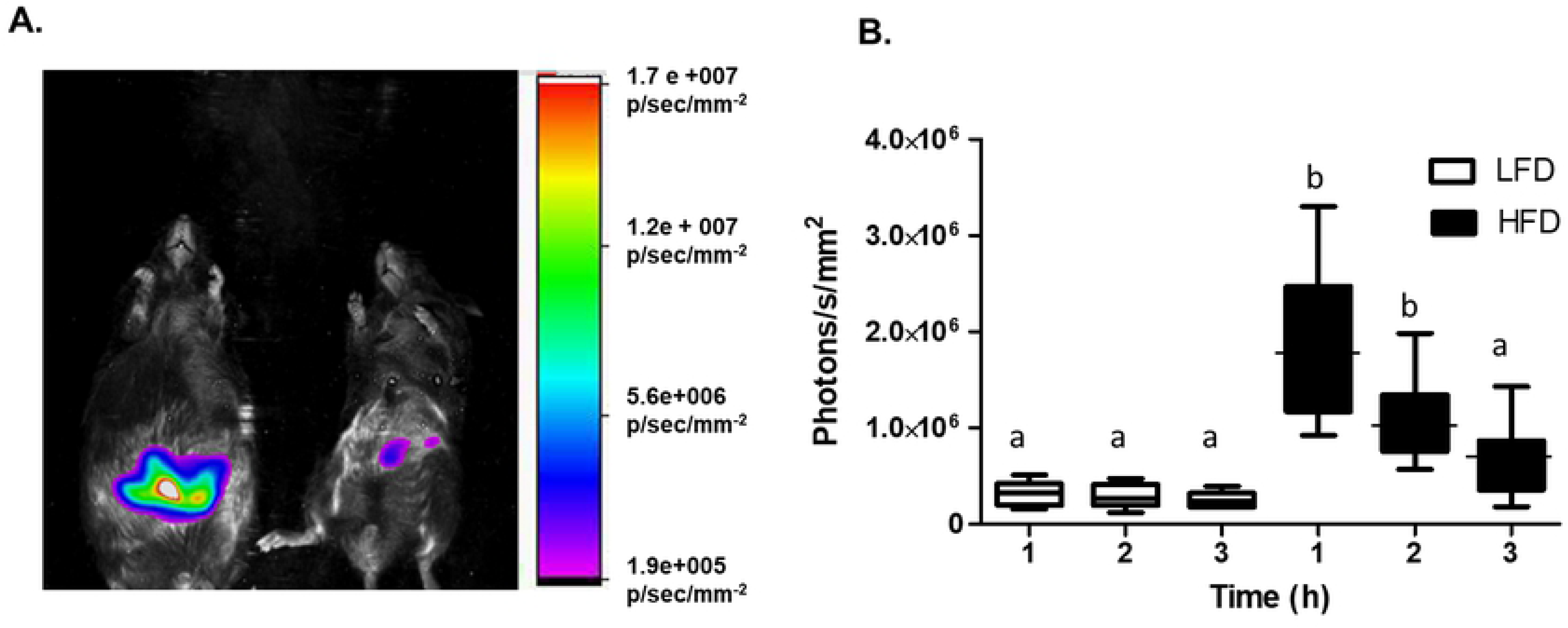
HFD-fed obese mice have higher levels of intestinal ROS than LFD-fed lean mice. Mice fed HFD or LFD for 10 weeks (n = 6/ group) were imaged in stationary, ventral orientation (abdomen facing camera) 3 h after H-ICG administration (i.e. 4 h after GPE administration).**A.** Representative overlay of ROS-associated NIRF image and corresponding brightfield image of a HFD-fed obese mouse (left) and a LFD-fed lean mouse (right). NIRF intensity scale shown on the right, image was normalized accordingly using Carestream MI software. **B.** ROS-associated NIRF in healthy, lean mice (white bars) and in obese mice (black bars) at 1, 2, and 3 h after administration of H-ICG. N=6 mice per group; Data are reported as mean ± SD. One-way ANOVA followed by the Tukey’s multiple comparison test was performed across both groups and all time points. Same letters indicate no difference between groups or time points while different letters indicate significant difference (p < 0.05).

### Comparison of ROS content in HFD (obese) and LFD (normal) mice over 360 ° of rotation

For all subsequent experiments, the MARS rotational system was used to capture both brightfield and NIRF images of individual mice at precise and repeatable angles to maximize the detection of NIRF emitted by ROS-sensitive dye. Total recorded NIRF was integrated over the full 360 ° turn with NIRF images superimposed on the brightfield images taken at every 30° turn (**Fig. 3A**). Compared to LFD-fed mice, the intestinal tract of HFD-fed animals contained significantly higher levels of ROS (**Fig. 3**). Plots of NIRF intensity at every 30° turn (**Fig. 3B**) and calculation of total area under the NIRF curve (**Fig. 3C**) further confirmed the presence of significantly greater quantities of ROS, detected by H-ICG florescence, in the intestines of the HFD-fed, obese and hyperglycemic mice. Specifically, compared to LFD-fed mice, HFD-fed mice had 4.1-times greater ROS-associated NIRF using rotational imaging (1.1 × 10^8^vs. 2.7 × 10^7^[photons*deg]/s/mm^2^AUC; Mann Whitney test, p< 0.0001) and 3.8-times greater ROS-associated NIRF using a single recording in ventral orientation (6.9 × 10^5^vs. 1.8 × 10^5^photons/s/mm^2^; Mann Whitney test, p< 0.0001; **Fig. 3C**), which is denoted as 0° in Figure 3A. As expected for light emanating from the intestines, ventral orientation towards the camera produced the highest NIRF. It is possible that our measurements underestimate the differences between obese and lean animals as the NIRF emitted from obese mice had to pass through a thicker layer of adipose tissue.

**Figure 3.**
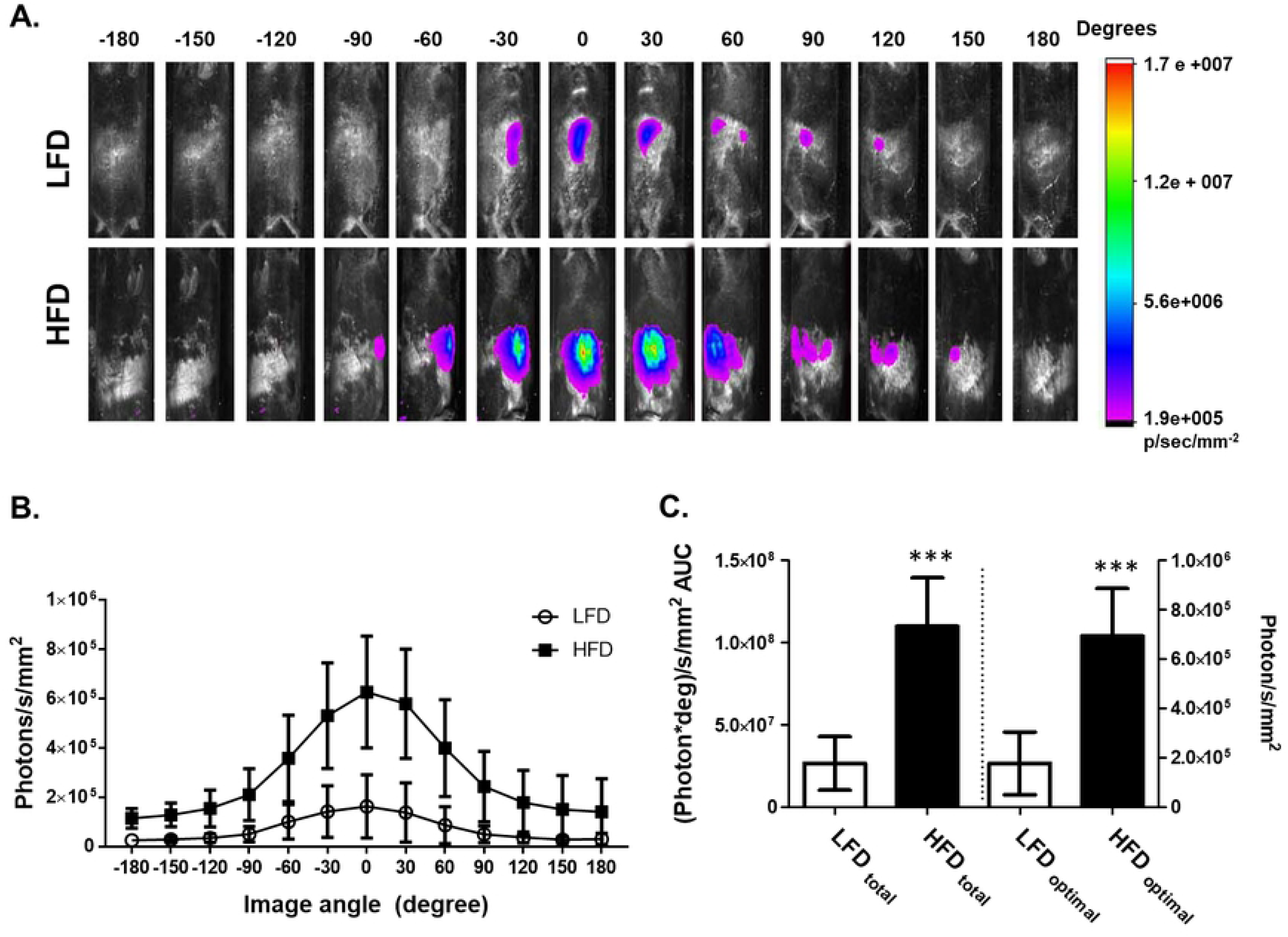
ROS-associated NIRF measured over a 360°rotational scan is higher in obese mice than in lean mice. **A.** ROS-associated NIRF images in obese (HFD) and lean (LFD) mice superimposed onto brightfield images. Rotation started 45 min following H-ICG administration, which was 1 h 45 min after GPE administration. Images were taken at 30° rotational increments over a course of a 360°rotation completed in 24 min. Images were colorized using Carestream MI software according to the NIRF intensity scale shown on right. **B.** Intensity of NIRF measured at different orientational angles in HFD and LFD-fed mice. Zero angle represents ventral orientation (abdomen facing camera). **C.** Area under curve (AUC) calculated from panel B (left axis) and NIRF measured at 0° corresponding to the ventral orientation (right axis). N= 20 mice per group; Data are reported as mean ± SD. Significant difference between HFD and LFD groups was detected using unpaired Mann Whitney test, *** p < 0.0001.

### Grape polyphenols reduce detectable ROS in the intestines of HFD and LFD-fed mice

We compared the ability of GPE, its major antioxidant constituents – PAC [46], and other dietary antioxidants, such as β-carotene, and a mixture of vitamins C, E and α-lipoic acid (ATL mixture), to reduce ROS in the mouse intestine. GPE, PAC and β-carotene represent relatively poorly bioavailable dietary antioxidants, while vitamins C, E and α-lipoic acid are all relatively bioavailable and relatively less stable in the digestive tract. Compared to obese, HFD-fed mice treated with vehicle (water), animals gavaged with GPE, PAC, or β-carotene had significantly reduced ROS in their intestines: 2.7-fold less for GPE and 1.9-fold less for both PAC and β-carotene (**Fig. 4A-B**). In contrast, the ATL mixture did not change the ROS-induced fluorescence (**Fig. 4A-B**).

**Figure 4.**
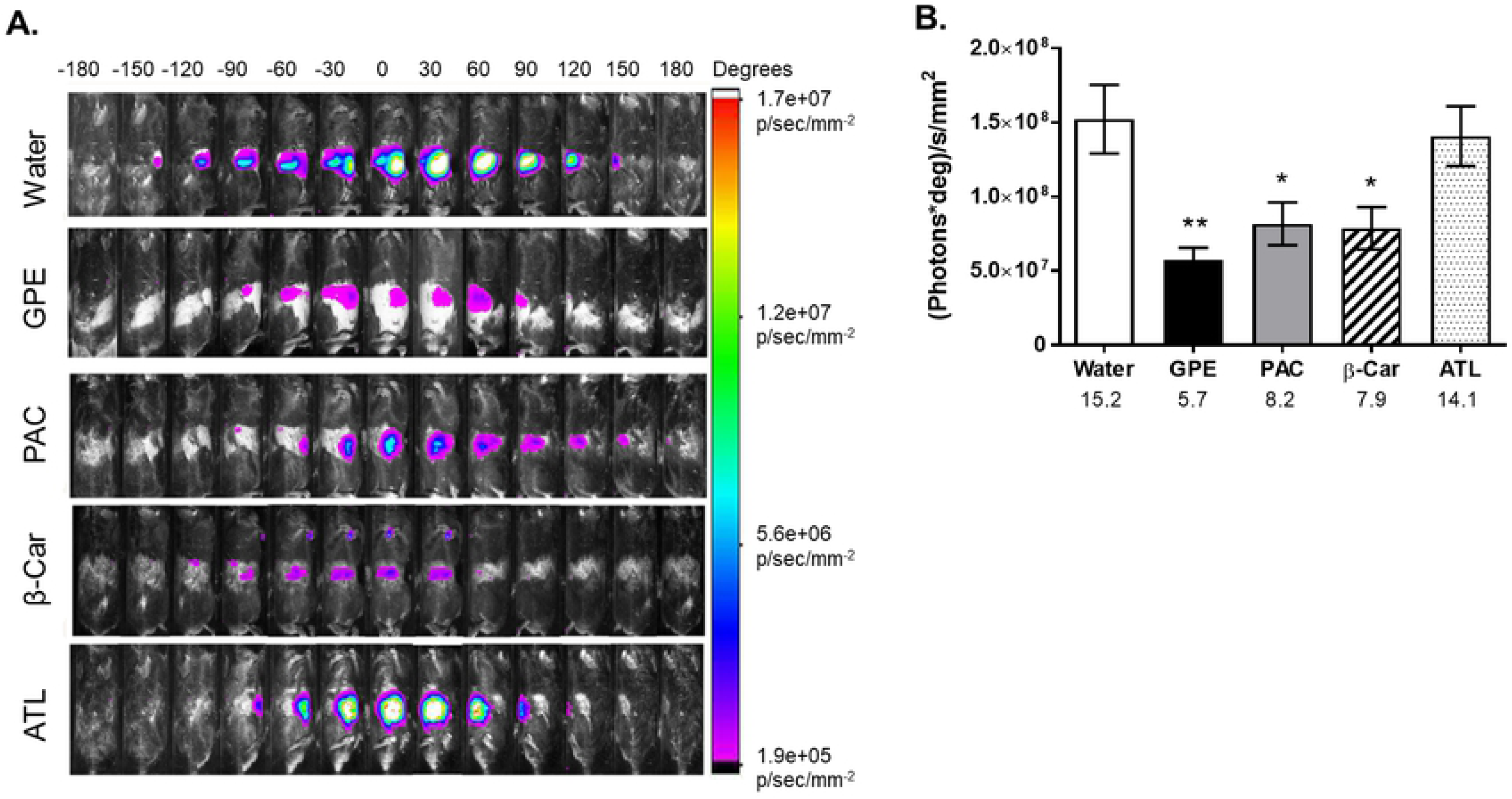
ROS-associated rotational NIRF of obese, HFD-fed mice treated with dietary antioxidants. **A.** ROS-associated 360°rotational NIRF images of obese HFD-fed mice superimposed on brightfield images. Animals were gavaged (1 h 45 min before imaging) with GPE, B-type proanthocyanidins (PAC), β-carotene (β-car), or ATL (mixture of L-ascorbic acid, D-α-tocopherol succinate and α-lipoic acid) at 32 mg/kg dose, except for GPE, which was dosed to deliver 32 mg/kg dose of total polyphenols. Images were taken at 30° rotational increments over a course of a 360°rotation completed in 24 min. Images were colorized using Carestream MI software according to the NIRF intensity scale shown on right. **B.** Area under curve (AUC) calculated from 4A as a function of different antioxidant treatments. Numbers under x-axis are x10^7^(photons*deg)/s/mm^2^AUC for the corresponding group. N= 5 mice per group. Data are reported as mean ± SD. Significant difference between groups was detected by one-way ANOVA (p= 0.002) followed by *post hoc* comparison to water-treated group using Dunnett’s test, * p< 0.05, ** p< 0.01.

Similar results were obtained in LFD-fed, lean mice subjected to the same treatments, although the LFD-fed mice consistently exhibited lower basal levels of ROS **(Fig. 5)**. Specifically, ROS-associated NIRF was reduced 2.3 times by GPE treatment, 2.4 times by PACs, 2.8 by β-carotene, and 1.9 times for ATL, although the latter was not significantly different from the control or other three treatments at p≤ 0.05.

**Figure 5.**
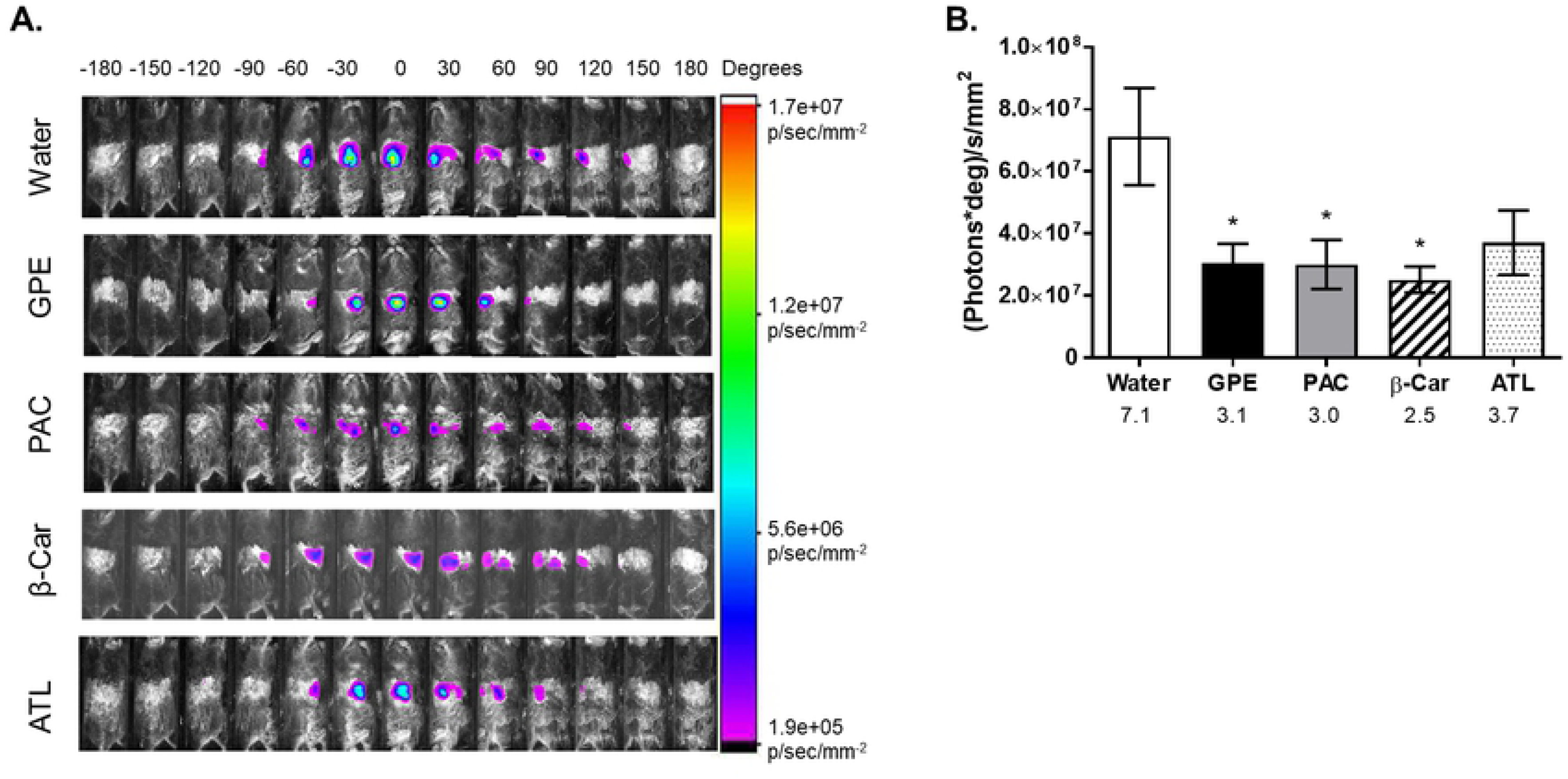
ROS-associated rotational NIRF of lean, LFD-fed mice treated with dietary antioxidants. **A.** ROS-associated 360° rotational NIRF images of obese LFD-fed mice superimposed on brightfield images. Animals were gavaged (1 h 45 min before imaging) with GPE, B-type proanthocyanidins (PAC), β-carotene (β-car), or ATL (mixture of L-ascorbic acid, D-α-tocopherol succinate and α-lipoic acid) at 32 mg/kg dose, except for GPE which was dosed to deliver 32 mg/kg dose of total polyphenols. Images were taken at 30° rotational increments over a course of a 360° rotation completed in 24 min. Images were colorized using Carestream MI software according to the NIRF intensity scale shown on right. **B.** Area under curve (AUC) calculated from 5A as a function of different antioxidant treatments. Numbers under x-axis are x10^7^(photons*deg)/s/mm^2^AUC for the corresponding group. N=5 mice per group; Data are reported as mean ± SD. Significant difference between groups was detected by one-way ANOVA (p= 0.023) followed by *post hoc* comparison to water-treated group using Dunnett’s test, * p< 0.05.

## DISCUSSION

Our data demonstrate that hyperglycemia and obesity in HFD-fed mice is associated with the higher content of ROS in the gut. This increased ROS content can be at least partially explained by the leaky gut syndrome usually associated with MetS and T2D [31, 50]. Increased permeability of the intestinal wall should allow more oxygen from mesenteric vasculature to diffuse into the lumen. Our data, for the first time, confirmed and visualized this phenomenon, as the increase in ROS in the gut of obese mice is likely associated with higher oxygen tension (**Fig. 2**). The data also showed that oral administration of poorly bioavailable dietary antioxidants such as GPE, its main component PAC, and β-carotene can reduce the content of ROS in the intestines of both obese and lean mice. In the case of obese mice, antioxidants, particularly grape polyphenols, reduced ROS content to the levels observed in the lean mice. This observation sheds light on the mechanism of action of dietary antioxidants to improve carbohydrate metabolism and prevent the development of T2D and obesity documented in multiple animal and human studies (see introduction). It may also provide some clues about the reason for earlier reported beneficial changes in gut microbiome and intestinal permeability associated with administration of poorly bioavailable dietary polyphenols [8-10].

It is tempting to speculate that very significant changes in gut redox potential associated with increased oxygen tension and ROS content in the gut will change the gut ecosystem, favoring allegedly beneficial anaerobic species, such as *A. muciniphila,* whose presence positively correlates with the improvements in symptoms of MetS and T2D [21-23, 51]. ROS are the most reactive and toxic forms of oxygen, particularly for anaerobic or microaerophilic organisms, such as *A. muciniphila,* which have no antioxidant systems to protect them [37]. Dietary polyphenols may, at least partially, act to reduce ROS and restore a more favorable redox potential in the gut thus promoting a healthier gut microbial environment. However, it is also possible that poorly bioavailable berry polyphenols exert their beneficial effects on health and microbiome through their antibiotic effects [52], by being partially metabolized by colonic bacteria to pharmacologically active compounds [20, 53], or by directly affecting growth of some gut bacteria as nutrients or regulators. However, our data suggest that close to 50% of the higher molecular weight polyphenols from GPE, which comprises mostly PAC, pass through the digestive system and can be recovered in the feces along with antioxidant activity they confer (**Fig. 1**). This observation downplays the possibility of GPE action through colonic metabolites or as nutritional substrates.

A mixture of more bioavailable dietary antioxidants comprising vitamins C, E, and α-lipoic acid was less effective in reducing intestinal ROS, particularly in the HFD-fed mice. We attribute this lack of activity to the fact that these essential nutrients are quickly absorbed or degraded during the intestinal transit, and thus cannot significantly affect ROS content and redox potential in the lower parts of the intestine.

Data presented in this manuscript may be one of the first demonstrations of how the antioxidant properties of dietary components are mechanistically linked to the prevention or mitigation of T2D and obesity and how these conditions are linked to the accumulation of ROS and to gut microbial ecology. We observed that dietary antioxidants, such as polyphenols in GPE, effectively mitigated HFD-induced accumulation of ROS in the gut and in the feces. Earlier, we have shown that GPE reduced endotoxemia, inflammatory cytokines, hyperglycemia and adiposity in HFD-fed mice while promoting a marked bloom of *A. muciniphila* in the gut [8].

In conclusion, this manuscript may begin to explain the mechanisms behind the health benefits of grapes and other fruits rich in poorly bioavailable antioxidants. The presented data suggest that beneficial effects of poorly bioavailable grape polyphenols (mostly PACs) on MetS/T2D, obesity, and hyperglycemia may be mediated through changes in gut microbiome triggered by their ROS scavenging activity. Specifically, reduction in ROS may benefit anaerobic and microaerophilic gut bacteria, such as *A. muciniphila*, that reside near the intestinal epithelium through which oxygen and ROS leak into the lumen. Well-documented bloom of *A. muciniphila,* which has been associated with anti-diabetic effects and production of anti-diabetic peptides such as Amuc_1100* [8, 9, 21-23], may be one of the events responsible for the reduction of hyperglycemia, reduced fat accumulation and improvements in molecular markers for intestinal health associated with dietary administration of GPE. Therefore, reducing the intestinal ROS associated with dysbiosis and MetS may be a promising target for developing new pharmaceutical or dietary therapies. Nevertheless, we cannot exclude the possibility that GPE-induced improvements in MetS are mediated by parallel, still undiscovered mechanism(s) that are independent of the antioxidant effects of GPE.

## Acknowledgments

IR is the guarantor for this manuscript. The authors thank Derek Adler for technical assistance with NIRF imaging and Kristin Moskal for help with preparing GPE.

## Figure legend to Supplementary Figure 1

**Supplementary Figure 1**. (+)ESI MS single ion chromatograms extracted at the corresponding m/z of individual proanthocyanidins (PACs) in GPE sample. **A** – oligomeric PACs; **B** – oligomeric PAC monogallates.

